# Determinants of Astrocytic Pathology in Stem Cell Models of Primary Tauopathies

**DOI:** 10.1101/2023.07.18.549558

**Authors:** Kimberly L. Fiock, Jordan Hook, Marco M. Hefti

**Author notes:** To whom correspondence should be addressed: Marco M. Hefti, MD, 25 S Grand Ave MRC-108-A Iowa City, IA 52240.

## Abstract

Astrocytic tau aggregates are seen in several primary and secondary tauopathies, including progressive supranuclear palsy (PSP), corticobasal degeneration (CBD), and chronic traumatic encephalopathy (CTE). In all cases, astrocytic tau consists exclusively of the longer (4R) tau isoform, even when adjacent neuronal aggregates consist of a mixture of 3- and 4R tau, as in CTE. The reasons for this and the mechanisms by which astrocytic tau aggregates form remain unclear. We used a combination of RNA *in situ* hybridization and immunofluorescence in post-mortem human brain tissue, as well as tau uptake studies in human stem cell-derived astrocytes, to determine the origins of astrocytic tau in 4R tauopathies. We found that astrocytes across tauopathies do not upregulate tau mRNA expression between diseases or between tau-positive and -negative astrocytes within PSP. We then found that stem cell-derived astrocytes preferentially take up long isoform (4R) labeled recombinant tau and that this uptake is impaired by induction of reactivity with inflammatory stimuli or nutritional stress. Astrocytes exposed to either 3R or 4R tau also showed downregulation of genes related to astrocyte differentiation. Our findings suggest that astrocytes preferentially take up neuronal 4R tau from the extracellular space, which potentially explains why astrocytic tau aggregates contain only 4R tau, and that tau uptake is impaired by decreased nutrient availability or neuroinflammation, both of which are common in the aging brain.

## INTRODUCTION

Neurodegenerative tauopathies can be classified based on the composition of pathological tau aggregates into pure short (3R) isoform, pure long (4R) isoform, or a mixture of both. Interestingly, only 4R tauopathies, such as corticobasal degeneration (CBD) or progressive supranuclear palsy (PSP), develop astrocytic tau pathology (**Fig. 1**). Tau pathology in Pick’s disease, a pure 3R tauopathy, and mixed tauopathies such as Alzheimer’s disease (AD) and primary age-related tauopathy (PART) is limited to neurons (**Fig. 1**). Even in chronic traumatic encephalopathy (CTE), where the pathognomonic neuronal tau aggregates in the depths of sulci are composed of a mixture of 3R and 4R tau, astrocytic tau aggregates are pure 4R [4]. In rodent models, astrocytic tau cannot propagate in the absence of neuronal tau expression [24], and human single-cell sequencing and RNA *in situ* hybridization data do not show an upregulation of tau expression in astrocytes from PSP patients [13, 25]. The human studies are, however, based on small numbers of cases, and CBD, which also has astrocytic tau aggregates, has not been systematically studied.

**Figure 1.**
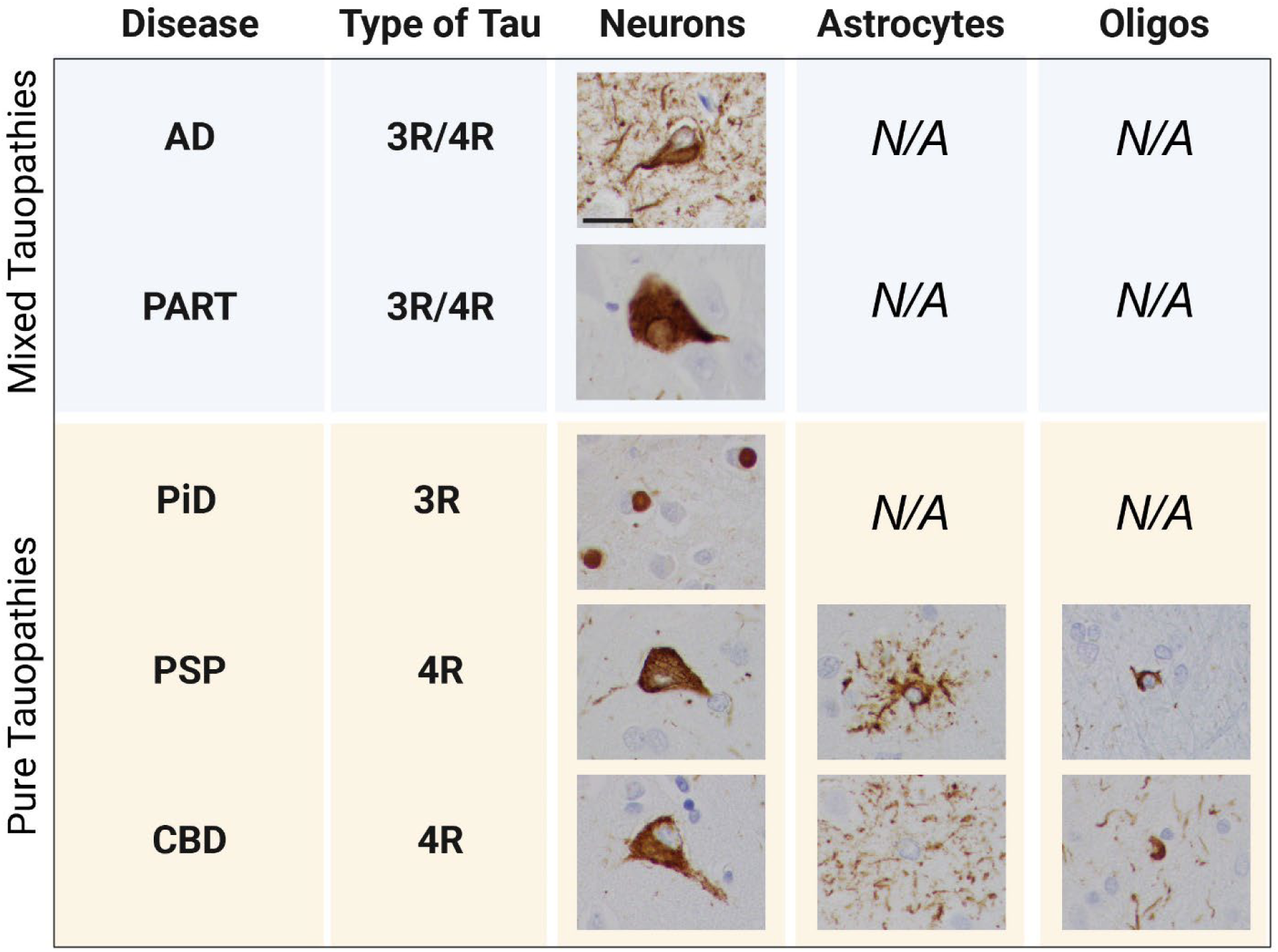
Tau pathology across diseases. Tau pathology is found exclusively in neurons in mixed tauopathies (AD and PART), but in neurons, astrocytes, and oligodendrocytes in pure tauopathies (PSP and CBD). PiD, a pure tauopathy, has predominantly neuronal pathology with rare astrocytic involvement. Scale bar=20μm

In disease, astrocytes undergo morphologic and functional changes in a process called reactive astrogliosis, which changes how they respond to and interact with their environment, including neurons and other astrocytes [27]. In stroke, for example, reactive astrocytes are responsible for glial scar formation, which sequesters damaged tissue to prevent further injury [28, 29]. Recently, the role of reactive astrocytes has been investigated in the context of neurodegenerative disease. Reactive astrocytes respond to pathologic cues by releasing extracellular signaling molecules which determine whether they play a neuroprotective or neurotoxic role [20, 35]. A1 reactive astrocytes are believed to release proinflammatory factors and cytokines that contribute to neuronal death while A2 reactive astrocytes are thought to release neurotrophic factors that can promote neuronal growth and survival [21]. While these astrocytic phenotypes have been addressed in diseases like AD, PD, and ALS, FTLD-tauopathies have largely been ignored or have been grouped in with other tauopathies under the umbrella term of “dementia” [18, 22]. This could be because a study done in 2002 suggested astrocytic tau pathology represents a degenerative process rather than a reactive one, though more recent work has shown upregulation of many of the genes associated with reactivity in humans with FTLD-tau and mouse models with familial FTLD-tau mutations [9, 16, 33].

Recent studies also suggest that astrocyte responses to disease are more complex than previously suspected and may require the use of multiple markers and assays to determine the presence of reactivity and its downstream impacts on functionality [8]. This indicates that definitive labels like “neurotoxic” or “neurotrophic” may not capture astrocytes’ true heterogeneity and that more work in this area is warranted to better understand how different stimuli contribute to various reactive phenotypes.

It has been well established that astrocytes readily take up both filamentous and monomeric recombinant tau *in vitro* [14, 17, 34], but the ability of astrocytes to take up different tau isoforms (e.g., 3R vs. 4R) remains largely unknown. Here, we report the first systematic assessment of astrocytic tau expression in human brain tissue across multiple neurodegenerative disorders and the novel findings that astrocytes preferentially take up 4R tau monomers, which is impaired by exposure to inflammatory stimuli or nutritional stress.

## RESULTS

### Astrocytic tau in human 4R tauopathies is of neuronal origin

We first sought to determine whether astrocytes in primary tauopathies, such as corticobasal degeneration and progressive supranuclear palsy, increase their expression of tau mRNA. Building upon our existing data using combined RNA *in situ* hybridization (RNAscope) and immunofluorescence, we found that the mean number of *MAPT* (tau) mRNA puncta per astrocyte (defined by positive immunofluorescence for GFAP) (**Fig. 2a**) and the proportional of astrocytes with puncta (**Fig. 2b**) did not differ significantly between adult control, Alzheimer disease, PSP, and CBD cases [10]. We then sought to determine whether astrocytes containing tau aggregates (GFAP^+^AT8^+^) had higher levels of tau mRNA than those without (GFAP^+^AT8^-^). Comparing GFAP^+^AT8^+^ to GFAP^+^AT8^-^ astrocytes *within* cases with progressive supranuclear palsy (PSP), we found no statistically significant difference in the mean number of puncta per astrocyte (**Fig. 2c**) or in the number of astrocytes expressing tau mRNA (not shown).

**Figure 2.**
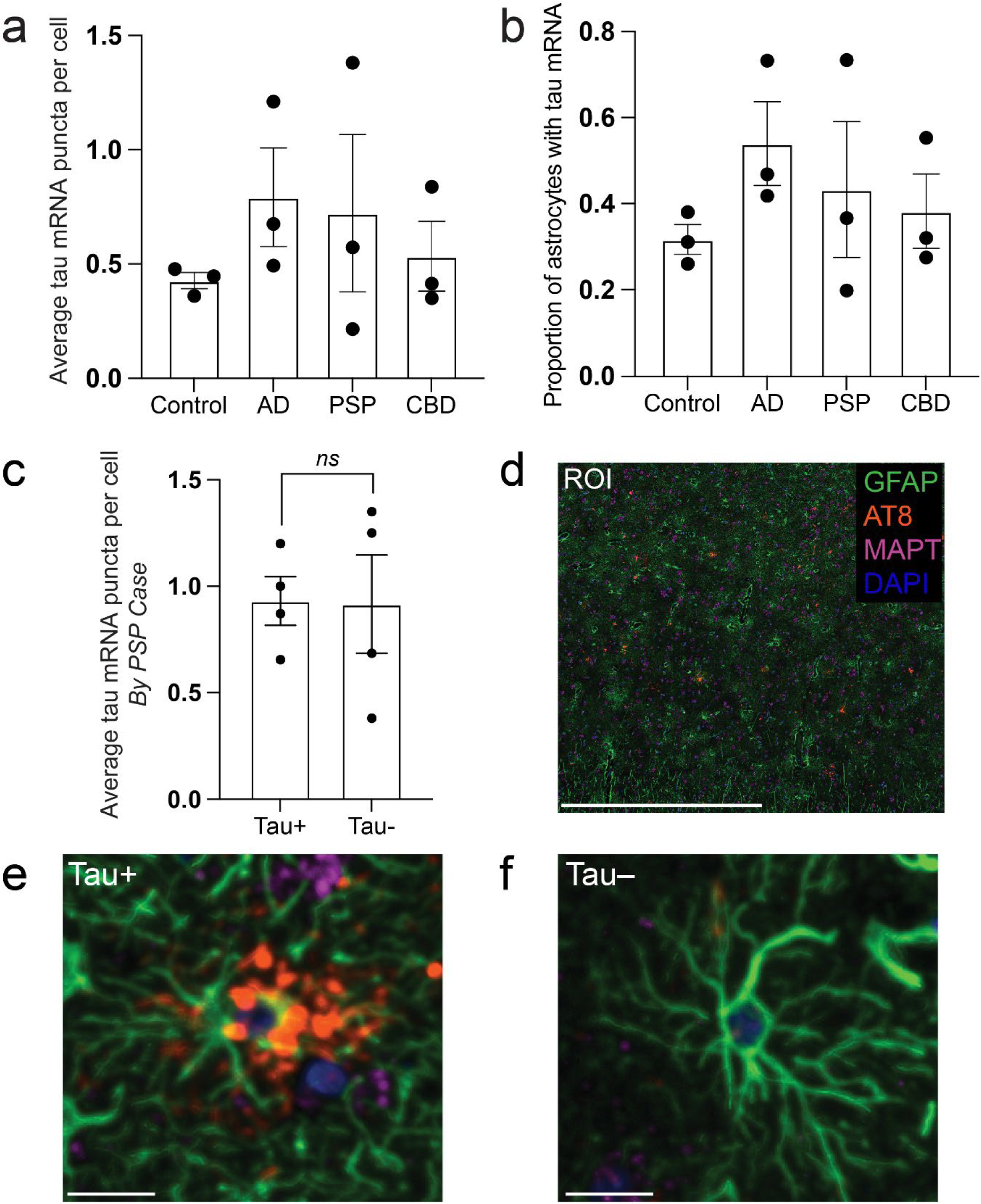
Astrocytes do not upregulate tau mRNA in astrocytic tauopathies. (A) Proportion of tau mRNA positive astrocytes in control, AD, PSP, and CBD. (B) Mean number of mRNA puncta per astrocyte in control, AD, PSP, and CBD. (C) Mean number of puncta comparing tau-positive and negative astrocytes within cases of PSP. (D) Representative low power ROI with insets of tau-positive and negative astrocytes shown in (E) and (F), respectively. Scale bars=1000 μm in (D) and 20 μm in (E) and (F); n=3 individual cases per condition in (A) and (B) and n=5 individual cases in (C)

### hESC-derived astrocytes preferentially take up 4R tau

Next, we wanted to characterize the ability of astrocytes to take up 3R and 4R tau using labeled recombinant (monomeric) tau and human stem cell-derived astrocytes. Stem cells and all subsequent differentiation steps were validated using a combination of immunocytochemistry (**Fig. 3, and S1-S3**) and trilineage differentiation (**Fig. S4**). Recombinant tau produced in *E. coli* was validated by SDS-PAGE, followed by Coomassie Blue staining and western blotting (**Fig. S5a** and **S5b,** respectively). When cultured with Cy5-labeled recombinant 1N3R or 1N4R tau, astrocytes took up significantly more 4R than 3R tau (p<10^-4^), with 4R uptake also occurring earlier than 3R (**Fig. 4**). This result was independently validated with a separate, independently differentiated, batch of astrocytes (**Fig. S6**).

**Figure 3.**
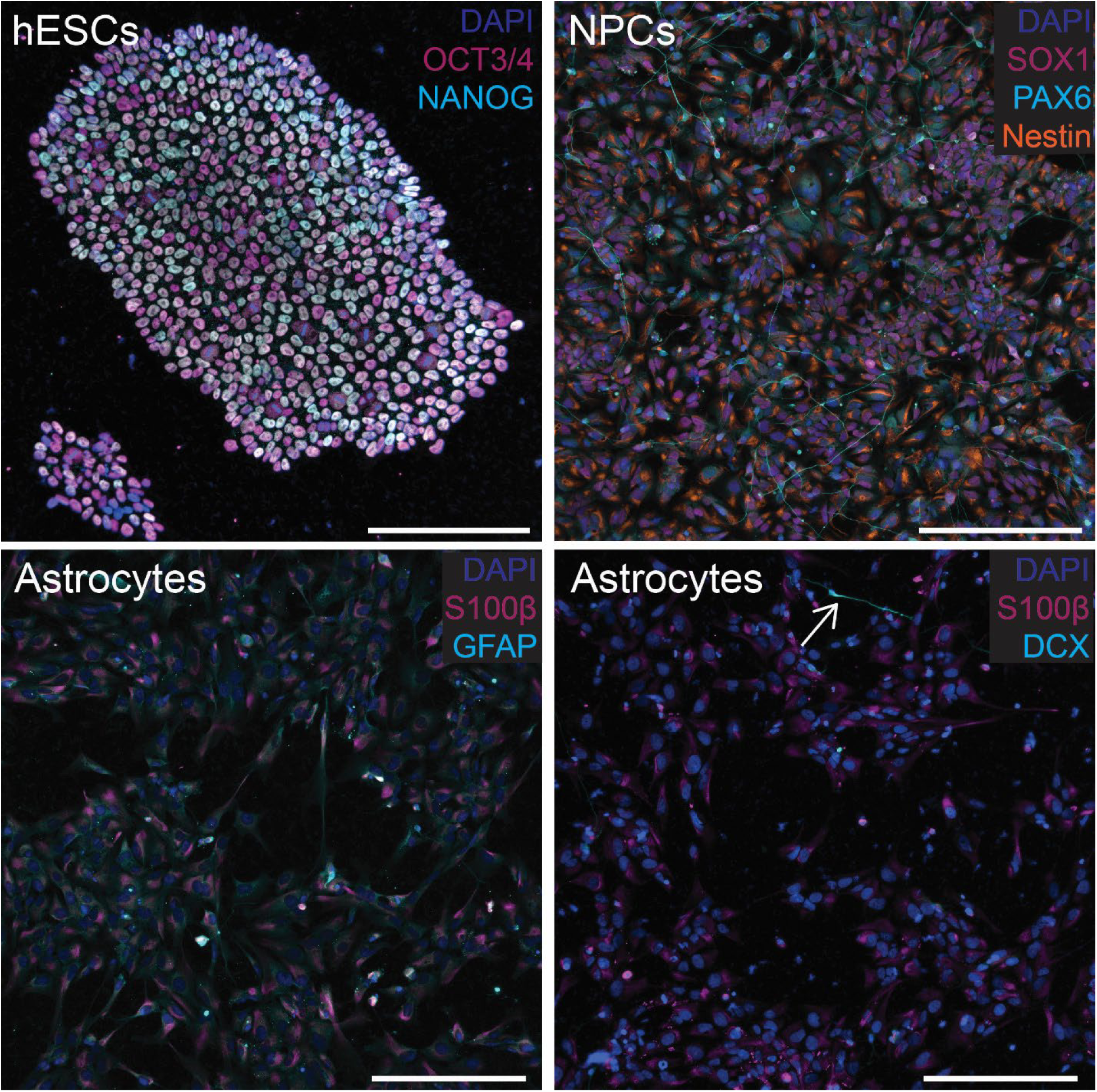
Validation of cell differentiation. (A) Stem cells show positive staining for OCT4 and NANOG, (B) neuronal precursor cells positive for Nestin, SOX1, and PAX6. (C) astrocyte staining for S100β and GFAP, (D) S100β and DCX. A single cell showing neuronal differentiation is indicated by white arrow in (D). Scale bars = 200 μm. Individuals panels for each stain are shown in Figs. S1, S2, and **S3** for stem cells, neural progenitors, and astrocytes, respectively

**Figure 4.**
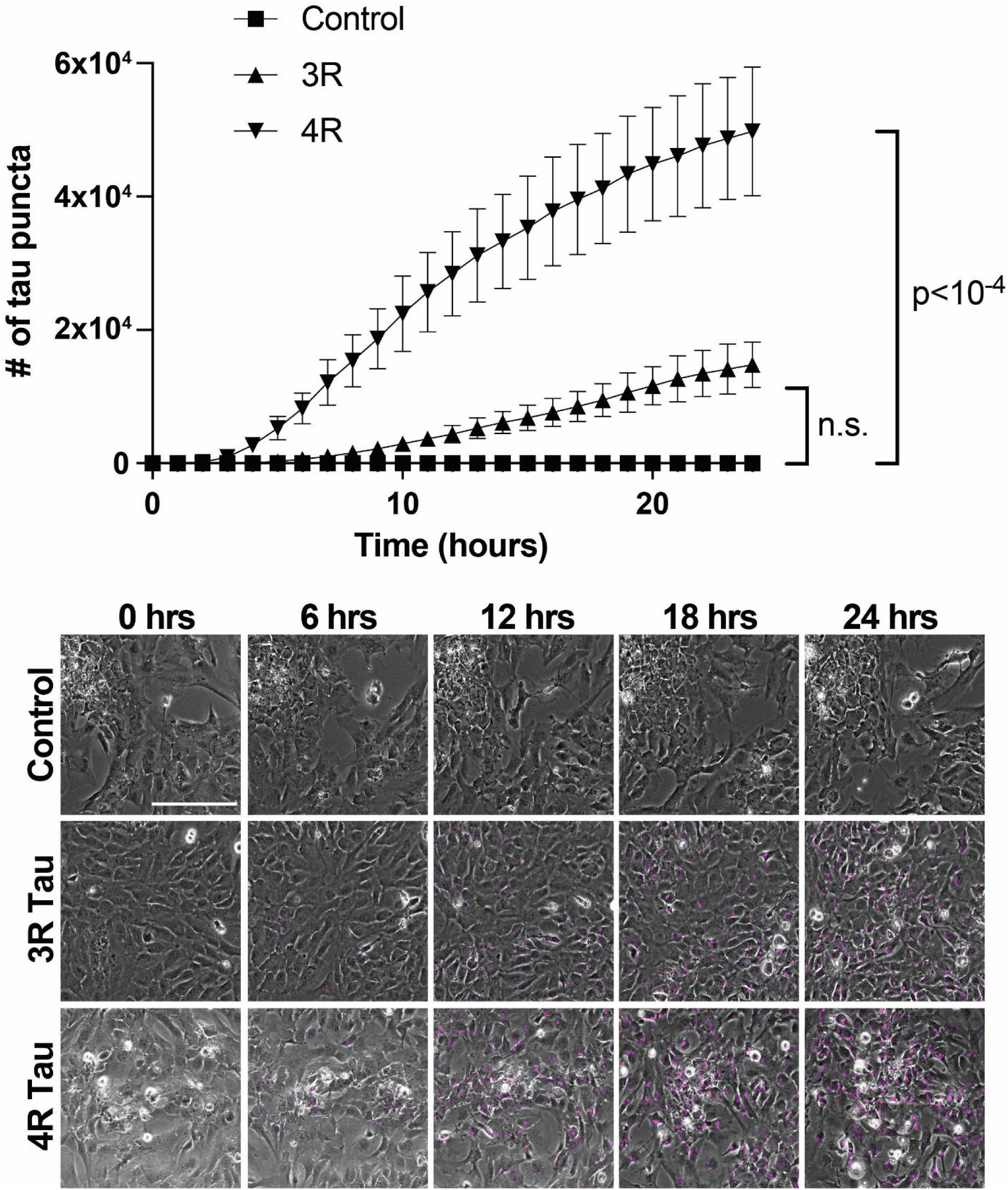
hESC-derived astrocytes preferentially take up 4R tau. hESC-derived astrocytes exposed to either labeled recombinant 3R or 4R tau over the course of 24 hours exhibit preferential uptake of 4R. *p<10^-4^ Image stills of hESC-derived astrocytes exposed to labeled recombinant 3R or 4R tau (magenta) over the course of 24 hours show more 4R in cells than 3R. Scale bar = 200μm; n=3 technical replicates per condition

### Astrocytic uptake of tau impairs astrocyte maturation

We then used bulk RNA sequencing to characterize the effect of 3R and 4R monomer uptake on astrocytes. We found 60 genes that differed between 3R and control, 29 between 4R and control, and no differentially expressed genes between 3R and 4R. Nine genes were significantly downregulated in both the 3R and 4R-exposed conditions compared to controls (**Fig. 5**). Five genes showed significant downregulation by at least a factor of two in both comparisons (C4orf48, HES4, INAFM1, TMEM59L, CAMK2N2). Of these down-regulated genes, C4orf48, HES4, INAFM1, and TMEM59L promote stem cell or astrocyte differentiation, which suggests that tau uptake impairs astrocyte differentiation [1, 7, 12].

**Figure 5.**
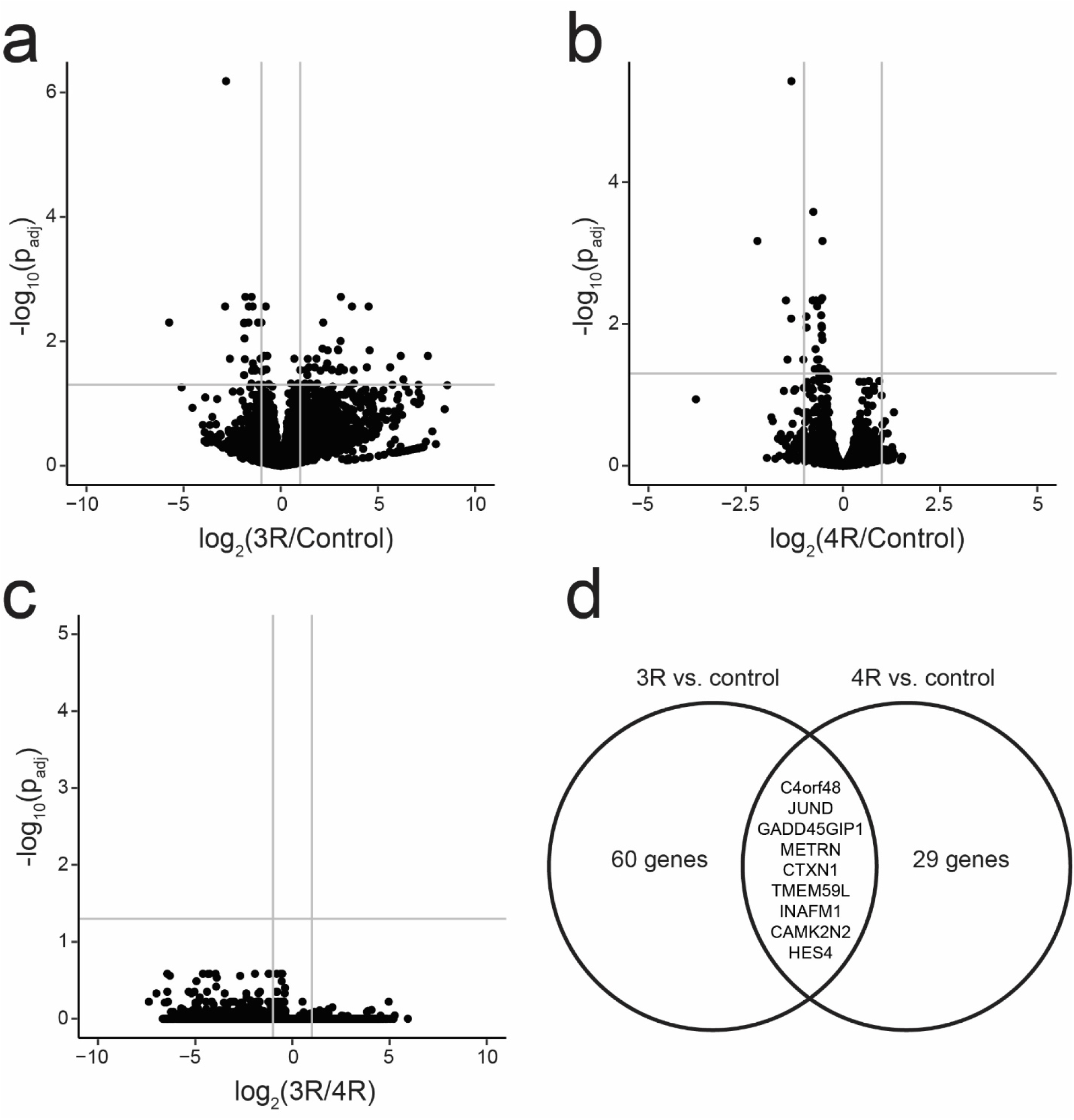
Effect of 3R and 4R tau uptake on astrocyte transcriptome. Volcano plots showing differentially expressed genes between 3R and control (A), 4R and control (B), and 3R and 4R (C). Genes differentially expressed in both 3R and 4R are shown in (D). Grey lines in (A) – (C) indicate cutoffs of adjusted p-value > 0.05 (horizontal) and absolute fold change > 2

### Astrocytes exposed to inflammatory stimuli or nutritional stress have impaired tau uptake and degradation

We then sought to test the effect of two distinct models of astrocytic stress on astrocytic tau uptake and degradation. We chose to focus on serum starvation, a model of nutritional stress, and an A1 inflammatory reactive astrocyte model using TNF-alpha, IL1-alpha, and C1q (“TIC”) [22]. The two models showed identical results, with both taking up significantly less 4R tau than their control counterparts (TIC vs. control, p=0.0015 and serum starved vs. control, p=0.0005) (**Fig. 6**). The amount of tau taken up between reactive astrocyte conditions, however, was not significantly different (**Fig. 6**). Interestingly, both conditions also reached a plateau in the amount of 4R taken up after roughly 10 hours, compared to the control condition that saw continuous uptake over the 24-hour period (**Fig. 6**). This suggests a possible impairment of degradation in the stressed astrocytes, where they cannot take up more tau due to a lack of clearance of the tau they already have.

**Figure 6.**
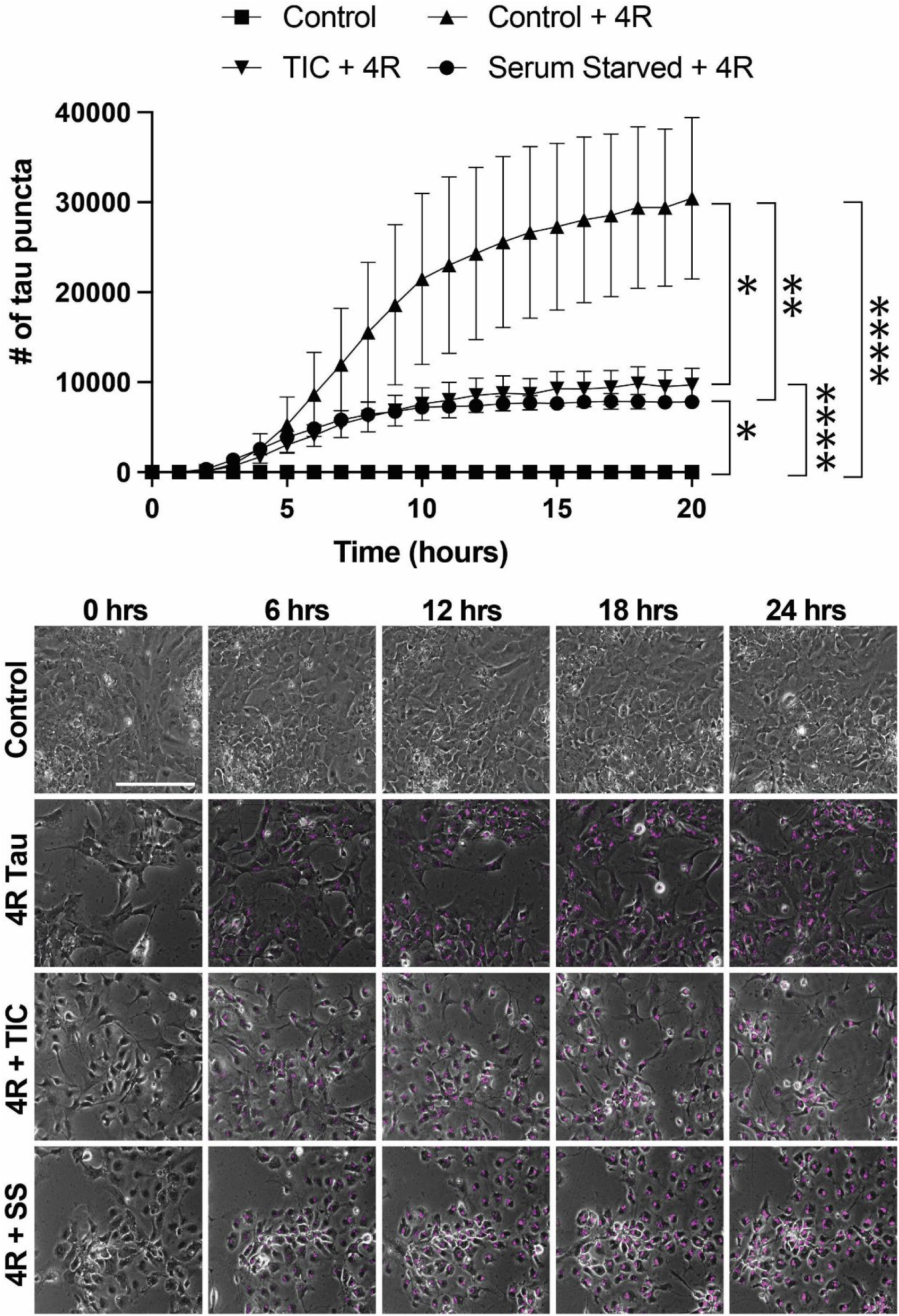
Nutritional or inflammatory stress impairs astrocyte tau uptake. hESC-derived astrocytes exposed to labeled 4R tau with or without serum starvation (SS) or TIC protocol (TNFα, IL1-α, C1q) with representative still images. N=3 technical replicates per condition; *p=0.0015, **p=0.0005, ***p=0.0001, ****p<10-5

## DISCUSSION

Our data shows that astrocytes do not increase tau expression in progressive supranuclear palsy or corticobasal degeneration. This is true *between* cases, when PSP or CBD are compared to controls, and *within* cases, comparing tau-positive and -negative astrocytes in patients with PSP. We then went on to show that stem cell-derived astrocytes preferentially take up 4R rather than 3R tau, that this downregulates the expression of genes related to astrocyte differentiation and is impaired by nutritional or inflammatory stress.

Our findings are broadly in agreement with single nuclear sequencing and RNA *in situ* hybridization studies in progressive supranuclear palsy [13, 25]. Our work is also complementary to and builds on data presented by Forrest et al in that we use GFAP immunofluorescence to better identify astrocytic mRNAs and include a larger number of cases. We also examined astrocytic tau expression in AD and CBD, which have not been previously characterized in histologic sections. Our findings are also complementary to work by other groups showing that astrocytes take up both tau monomers and oligomers [14, 17, 34]. Our research supports and builds on this data by demonstrating isoform-specific uptake of tau monomers, which has, to our knowledge, not been previously described.

Our studies are limited by the low throughput of our RNA *in situ* assays and downstream analytic pipeline, which limits the number of cases examined, and by the small number of channels available in our RNAscope workflow. We were also not able to determine whether there is a difference in tau expression between tau-positive and -negative astrocytes in corticobasal degeneration. This is due to the more diffuse nature of astrocytic plaques compared to the tufted astrocytes seen in PSP and the resulting difficulty in identifying the parent astrocytes in our thin (5 μm) sections. Answering this question will require three-dimensional reconstruction in thicker sections or the application of tissue-clearing methods. Novel spatial transcriptomic methods combined with protein stains to identify tau-positive astrocytes also show great promise for in-depth astrocytic phenotyping in PSP and CBD. Our uptake assays are inherently limited due to their reliance on recombinant tau monomers, which do not necessarily recapitulate the post-translational modification patterns seen in the human brain. The relative toxicity of tau monomers and oligomers, and their respective contributions to astrocytic tau pathology, likewise remains unknown.

In summary, we have shown that astrocytes in PSP and CBD do not upregulate tau expression, that they preferentially take up 4R tau monomers *in vitro*, and that this process is impaired by nutritional and inflammatory stress. These findings strongly suggest that the increased levels of astrocytic tau seen in PSP and CBD are due to neuronal uptake rather than upregulation of astrocytic tau production and identify a potential mechanism for the isoform-selectivity of astrocytic tau aggregates. Future studies will be necessary to determine the molecular mechanisms for this uptake and its downstream effects on the recipient astrocytes.

## METHODS

### Tissue procurement

Formalin-fixed paraffin-embedded (FFPE) tissue was obtained as previously described [10]. Individual cases used, demographic information, and sources are listed in **Table 1**. The University of Iowa’s Institutional Review Board determined that, since this project used tissue from deceased individuals exclusively, it does not represent human subjects research under the NIH common rule (determination #201706772). All methods were conducted in accordance with the relevant laws, regulations, guidelines, and ethical standards of our institution and with the 1964 Helsinki declaration and its later amendments or comparable ethical standards. Diagnoses were made according to the updated National Alzheimer’s Coordinating Center neuropathologic diagnostic criteria and other published guidelines [2, 5, 6, 15, 23].

**Table 1.** Demographic information and source of all human tissue samples used in current study.

### Combined RNA in situ hybridization and immunofluorescence

RNA *in situ* hybridization (RNAscope) with immunofluorescence (IF) was performed on formalin-fixed, paraffin embedded (FFPE) tissue using the 3-plex Multiplex Fluorescent v2 Reagent Kit (Cat No. 323100, ACDBio) with the C1-*MAPT* probe (Cat. No 408991, ACDBio) according to the manufacturer’s protocol and as previously described [10]. We have previously validated the *MAPT* probe using human tau transgenic mice [11].

### Slide scanning and quantification

Slide scanning and quantification were done using a Cytation 5 from Agilent (Cat No. BTCYT5PW) with GEN5PRIME software (Cat No. BTGEN5IPRIM, Agilent) to create a customized protocol. Each set of experiments was run using identical protocol settings. A discovery scan was done to image the entire piece of tissue at 4X (Cat No. BT1320515, Agilent) using a combination of the DAPI filter cube (Cube: Cat No. BT1225100, Agilent; LED: Cat No. BT1225007, Agilent), GFP filter cube (Cube: Cat No. BT1225101, Agilent; LED: Cat No. BT1225001, Agilent), TRITC filter cube (Cube: Cat No. BT1225125, Agilent; LED: Cat No. BT1225012, Agilent), and CY5 filter cube (Cube: Cat No. BT1225105, Agilent; LED: Cat No. BT1225005, Agilent) as appropriate. Five regions of interest (ROIs) were selected at random in the cortex for each tissue using approximately equal sized ROIs; then each ROI was imaged in the appropriate channels using the above filters at 40X (Cat No. BT1320518, Agilent) and stitched using the GFP channel as a reference with 150μm of overlap.

For quantification, subpopulations were defined in GEN5PRIME to identify cell types of interest and count the number of RNA puncta present. Astrocytes were defined as cells with a nucleus size of less than 15 based on DAPI and a mean GFP intensity of at least 10,000; astrocytes with tau pathology were defined as cells that met the above criteria in addition to a mean TRITC intensity of at least 10,000, and astrocytes without tau pathology were defined as cells that met the above criteria with a mean TRITC intensity of less than 10,000. RNA puncta were counted in each population based on the size of puncta (between 2-5μm) using either the CY5 channel alone or the TRITC and CY5 channels as appropriate. The average number of puncta in each subpopulation was calculated by GEN5PRIME using all five ROIs for each slide. P-values were assessed in GraphPad Prism using t-tests with each sample counting as one biological replicate, and graphs were made using R Studio and Adobe Illustrator. Data is presented as mean ± standard error of the mean (SEM).

### Stem cell procurement and culture

The H14 human embryonic stem cell (hESC) line was obtained from WiCell (Cat No. WAe014-A) and is listed as an NIH-approved line for research purposes (NIH Approval Number: NIHhESC-10-0064). Stem cell culture was done using a commercially available media kit (mTeSR Plus; Cat No. 100-0276, STEM CELL Technologies) following the manufacturer’s published protocol [32]. Briefly, cell colonies were grown on Matrigel-coated 6-well plates (Matrigel: Cat No. 354277, Corning; Plates: Cat No. CLS3516-50EA, Sigma Aldrich) in mTeSR Plus for at least three passages before being dissociated for downstream differentiation protocols. Matrigel was diluted according to the lot-specific dilution factor recommended by the manufacturer in 25 ml of DMEM/F12 HEPES (Cat No. 36254, STEMCELL Technologies) and set for 1 hour at 37°C before plates were used. Areas of cellular differentiation were removed each day from cultures. Passaging of colonies was done roughly every five days using Gentle Cell Dissociation Reagent (GCDR) from STEMCELL Technologies (Cat No. 100-0485). Stem cell pluripotency was validated using the STEMdiff Trilineage Differentiation Kit (STEMCELL Technologies, Cat No. 05230) and human stem cell antibody array from AbCam (Cat No. ab211066) according to the manufacturer’s directions.

### Generation of neural progenitor cells

Neural progenitor cells (NPCs) were created from the hESCs described above using the STEMdiff SMADi Neural Induction kit from STEMCELL Technologies (Cat No. 08581) according to their guidelines for Matrigel-coated 6-well plates [31]. Briefly, colonies were checked for areas of differentiation and removed, if necessary, before washing once with cold DPBS-/-. Colonies were then incubated for 8-10 minutes at 37°C in GCDR for dissociation. Cells were collected and triturated to create a single-cell suspension, then spun down at 300g for 5 minutes at room temperature. Cell counts were determined using a Countess 3 instrument (Cat No. AMQAX2000, Thermo Fisher), and 2,000,000 cells/well were plated in a 6-well plate in STEMdiff SMADi Neural Induction media. Daily media changes were performed for seven days followed by passaging of progenitor cells using Accutase (Cat No. 07920, STEMCELL Technologies), which was repeated for one additional passage (two passages total). Cells were plated at 1,500,000 cells/well in a 6-well plate for each passage. At passage 3, cells were either frozen down in CryoStor CS10 (Cat No. 07930, STEMCELL Technologies) at 3,000,000 cells/vial or moved onto the next step of the differentiation process.

### Generation of astrocyte precursor cells and mature astrocytes

Astrocyte precursor cells were created from the neural progenitor cells described above using the STEMdiff Astrocyte Differentiation media kit from STEMCELL Technologies (Cat No. 100-0013) following their protocol for Matrigel-coated 6-well plates [30]. Briefly, progenitor cells were plated at a seeding density of 1,900,000 cells/well in a Matrigel-coated 6-well plate and given daily media changes for seven days. For passaging, cells were incubated with Accutase for 8-10 minutes at 37°C then spun down at 400g for 5 minutes at room temperature. Cell counts were obtained using a Countess 3 instrument, and a density of 1,400,000 cells/well was used. Media changes were performed every other day for seven days, then the process was repeated once more (seeding density: 1,600,000 cells/well) for a total of two passages in the differentiation media. At passage 3, precursor cells were either frozen in CryoStor CS10 (above) at 2,000,000 cells/vial or moved onto the final stage of differentiation.

Mature astrocytes were created from astrocyte precursor cells using the STEMdiff Astrocyte Maturation media kit from STEMCELL Technologies (Cat No. 100-0016) following their guidelines for Matrigel-coated 6-well plates. Briefly, precursor cells were seeded at 1,900,000 cells/well in a 6-well plate, and media changes were performed every other day for seven days. Cells were dissociated with Accutase for 8-10 minutes at 37°C before being spun down at 400g for 5 minutes at room temperature. This process was repeated once more for 2 passages in the maturation media before cells were plated for downstream analysis.

### Immunocytochemistry

Primary and secondary antibodies used with their respective concentrations are shown in **Table 2**. Cells for ICC were grown in a 12-well plate (Cat No. 3513, Costar) on autoclave-sterilized 18mm coverslips (Cat No. 12545100, Fisher Scientific) coated with Matrigel using the same protocol as described above. Coverslips were washed once with 1X PBS before fixing for 30 minutes with 10% formalin at room temperature and were subsequently washed twice with 1X PBS before storing in 1X PBS+ 0.01% sodium azide at 4°C. Coverslips were placed in a 12-well plate and incubated for 10 minutes at room temperature in extraction solution (0.02% Triton-X 100 in 1XPBS; Triton-X 100: Cat No. BP151, Fisher Scientific) before blocking for 30 minutes at room temperature in blocking solution (10% heat-inactivated fetal bovine serum (hiFBS) in 1XPBS; hiFBS: Cat No. S1245914, R&D Systems). Primary antibody incubation was done with the respective antibodies diluted in serum solution (1% hiFBS in 1XPBS) overnight at 4°C with gentle rocking. Coverslips were washed five times for 1 minute each at room temperature with 1X PBS, then incubated in secondary antibody diluted in serum solution for 1 hour at room temperature protected from light with gentle rocking. Five 1-minute 1X PBS washes were performed before incubation with TrueView Autofluorescence Quenching solution (prepared according to manufacturer’s guidelines and diluted 1:10 with ddH_2_O) for 5 minutes with gentle agitation at room temperature protected from light. Coverslips were washed once more for 5 minutes with 1X PBS at room temperature with the addition of three drops of DAPI (Cat No. R37606, Invitrogen) then mounted on SuperFrost Plus slides (Cat No. 12-550-15, Fisher Scientific) using VECTASHIELD Plus mounting medium (Cat No. H-1900, Vector Laboratories).

**Table 2.** Antibodies used in study.

### Generation of plasmids

Custom human TauB and TauE plasmids were created on a pET29b backbone by GenScript on a fee-for-service basis, with 6xHIS and S tags added to the N- and C-terminal ends to ensure high purity of the resulting protein preparations. The resulting plasmids were validated by long-read sequencing (Plasmidsaurus), transformed into BL21 (DE3) OneShot Competent Cells (Cat No. C6000-03, Life Technologies), and given to the Carver College of Medicine’s Protein and Crystallography Facility at the University of Iowa for recombinant protein generation.

### Generation of recombinant tau

Recombinant human TauB (1N3R) and TauE (1N4R) proteins were produced at the Carver College of Medicine’s Protein and Crystallography Facility at the University of Iowa. Protein was produced in *E. coli* BL21 (DE3) cells (Cat No. 69450, Novagen) with the pET29b plasmids generated above. Bacteria were grown in Luria Broth (Cat No.910-79-40-2, Research Products International) in the presence of kanamycin (100 ug/mL), and protein overexpression was induced with 1 mM IPTG in 3 hours at 37°C. Cells were disrupted by sonication and boiled at 75°C for 15 minutes in buffer (50 mM NaPO_4_, 2 mM EDTA and 2 mM DTT, pH 6.8). All subsequent procedures were performed at 4°C. Lysate was ultracentrifuged at 80,000g for 45 minutes, and protein was purified by cation exchange chromatography (HiTrap SP, GE Healthcare), followed by size exclusion chromatography (HiLoad 16/600 Superdex 200 pg, GE Healthcare). Recombinant tau was labeled using an Alexa Fluor 647 Protein Labeling Kit (Cat No. A20173, Invitrogen) with 500 mg of recombinant protein input at a concentration of 1 mg/mL according to the manufacturer’s recommendations. Successful labeling was confirmed by measuring absorbance on a NanoDrop spectrophotometer.

### Western blotting

Western blots were run using 20ng of purified protein or 30μg of cell lysate. Samples were run on a 10% Mini-PROTEAN TGX Stain-Free Precast gel (Cat No. 4561036, Bio-Rad) and blotted to PVDF. The membrane was blocked in EveryBlot blocking buffer (Cat No. 12010020, Bio-Rad) for 10 minutes at room temperature, then HT7 was diluted 1:5000 in EveryBlot and incubated with the membrane overnight at 4°C (**Table 2**). Horseradish peroxidase-labeled horse anti-mouse secondary was used at 1:5000 diluted in EveryBlot for 1 hour at room temperature and detected by Clarity Western ECL substrate (Cat. No 1705060, Bio-Rad) (**Table 2**). Chemiluminescence was measured using a ChemiDoc Touch Imaging System (Cat No. 1708371, Bio-Rad).

### Astrocyte plating for live cell imaging

Mature astrocytes were plated in glass bottom 12-well plates (Cat No. P12-1.5H-N, Cellvis) at a density of 200,000 cells/well and allowed to settle for a minimum of 24 hours before live-cell imaging. For experiments requiring different media conditions, cells were plated in either STEMDiff Astrocyte Maturation media without Supplement B (serum-starved) or STEMDiff Astrocyte Maturation media without Supplement B with the addition of “TIC” proinflammatory factors (30ng/mL TNF-alpha (Cat No. 300-01A-10ug, Peprotech), 3ng/mL IL1-alpha (Cat No. 500-P21A-50ug, Peprotech), 400ng/mL C1q (Cat No. ab282858, Abcam) (TIC) [19]. Serum starvation occurred either at the beginning of the astrocyte maturation protocol (three weeks total) or for four days before live-cell imaging, and “TIC” induction occurred 24 hours prior to live-cell imaging.

### Live cell imaging

Live-cell imaging was done using a Cytation 5 (above) with GEN5PRIME software (above) for either 24- or 48 hours using phase contrast with laser autofocus (Cat No. BT1225010, Agilent) and the CY5 filter cube (above) at 20X (Cat No. BT1320517, Agilent). Cells were incubated in the imager at 37°C with 5% CO_2_ to mimic normal incubator conditions. A 5x5 matrix was used to capture 25 tiles across the center of the well with images captured every hour throughout the experiment. Images were processed in GEN5 to remove background fluorescence in the CY5 channel caused by phenol red in the media then tau puncta spots were counted by GEN5 using a minimum and maximum size cutoff (2.5μm and 10μm, respectively). Due to difficulties counting cells on phase contrast by GEN5, all phase contrast images for timepoint 1 (baseline) in each experiment were imported into Labscope (Zeiss). Cells were counted using the default settings of the AI Cell Counting Module (Zeiss), and counts were exported into a spreadsheet for data analysis.

### RNA sequencing

RNA extraction was performed from the mature hESC-derived astrocytes grown as a monolayer and used for live-cell imaging. Invitrogen’s PureLink RNA Mini kit (Cat No. 12183018A) was used for RNA extraction according to the manufacturer’s protocol, and samples were sent to GeneWhiz for sequencing. For samples sent as cell pellets, cells were grown as described above, then scraped into cold DPBS-/-(above) using a cell spatula and spun down at 400g for 5 minutes at 4°C. The supernatant was discarded, and pellets were flash-frozen in a dry ice/ethanol slurry. RNA extraction and sequencing were done by GeneWhiz’s sequencing facility on a fee-for-service basis. Data analysis was conducted as previously described [26]. Briefly, RNA sequencing reads were aligned to the hg38 human reference genome using STAR aligner. Reads were quantified using the *subreads* package, and comparative gene expression analysis was done using the *DESeq2* package in R. Gene ontology enrichment analysis was done using the web interface from the Gene Ontology Consortium with the biological process as the readout on all significant genes (p<0.05) [3].

### Statistical analysis

Data analysis for RNA sequencing is described above. For other studies, data analysis was done by either a generalized linear model with a HAC correction for time dependence or using paired or unpaired t-tests, as appropriate. Total cell count, treatment group, and position in well were also included as independent variables for the generalized linear model. Generalized linear models were run in MATLAB, t-tests were done in GraphPad Prism, and graphs were generated using either R studio or GraphPad Prism.

### Figure creation

All figures were made using Adobe Illustrator or the BioRender imaging platform.

## DATA AVAILABILITY STATEMENT

All data and code generated over the course of this study are provided in the supplemental material.

## ACKNOWLEDGEMENTS

The authors wish to thank histotechnologists Mariah Leidinger and Melissa Jans for their technical assistance with block preparation and sectioning, as well as decedent care specialists Adam Ciha, Benjamin Palmer, Terry Anderson, Michael Hanes, and Vaughn Dohse for their assistance with tissue procurement and autopsies. We thank the Brain Bank for Neurodegenerative Disorders at Mayo Clinic, Jacksonville, FL and the Iowa Neuropathology Resource Laboratory for providing us with tissue sections. The authors also wish to thank the donors and their families who made the work possible. This work was supported by grants from the Histochemical Society (to KLF), the NIH (K23 NS109284 to MMH), the Graduate College of the University of Iowa (to KLF), and the Roy J. Carver Foundation (to MMH).

## DECLARATIONS

This work was supported by grants from the Histochemical Society (to KLF), the NIH (K23 NS109284 to MMH), the Graduate College of the University of Iowa (to KLF), and the Roy J. Carver Foundation (to MMH). The authors have no other relevant disclosures.

