## Supplemental Data for "Determinants of Astrocytic Pathology in Stem Cell Models of Primary Tauopathies"

*To whom correspondence should be addressed:

Marco M. Hefti, MD

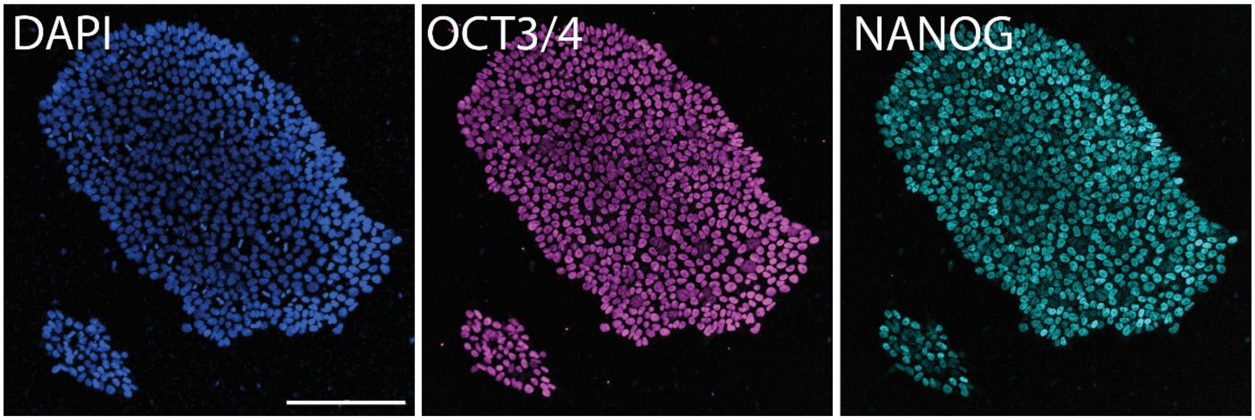


**Figure S1. Validation of stem cells by immunocytochemistry.** Channels shown separately for image in **Fig. 3**. Scale bar = 200 μm

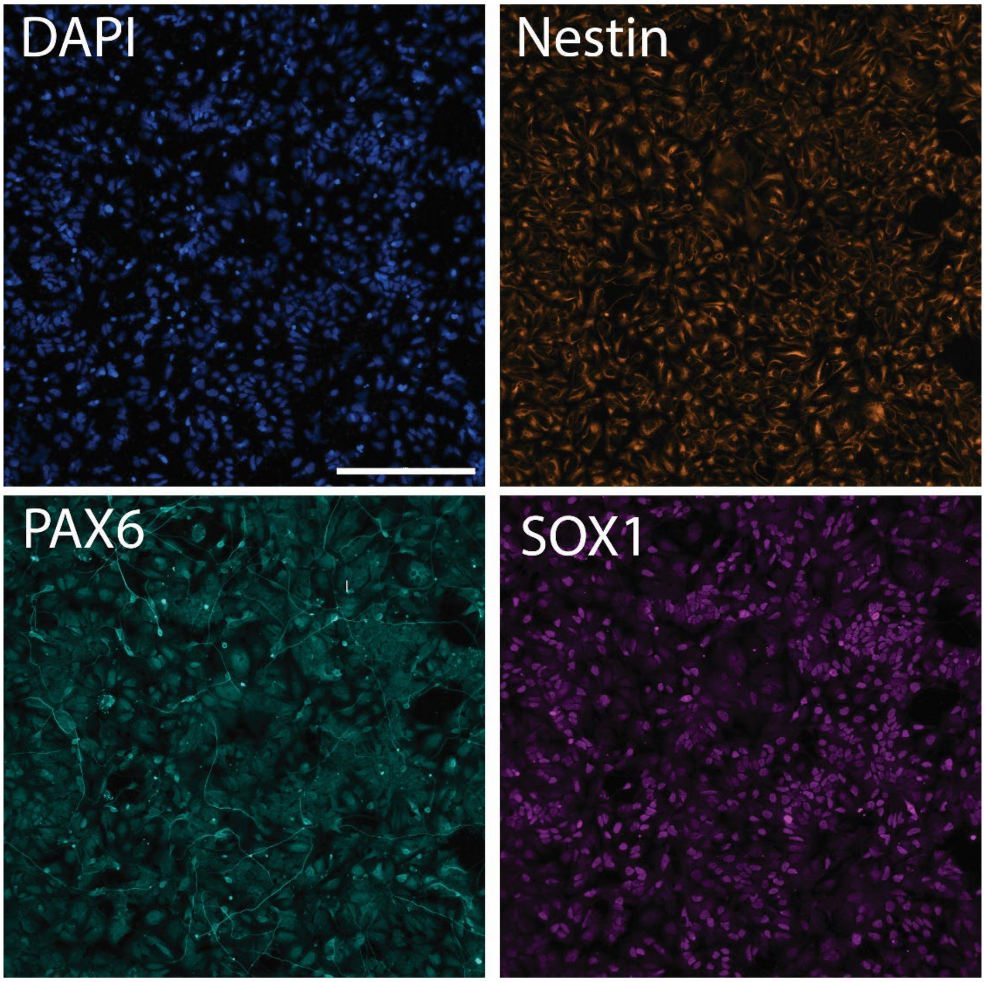


**Figure S2. Validation of neural progenitor cells by immunocytochemistry**. Channels shown separately for image in **Fig. 3**. Scale bar = 200 μm

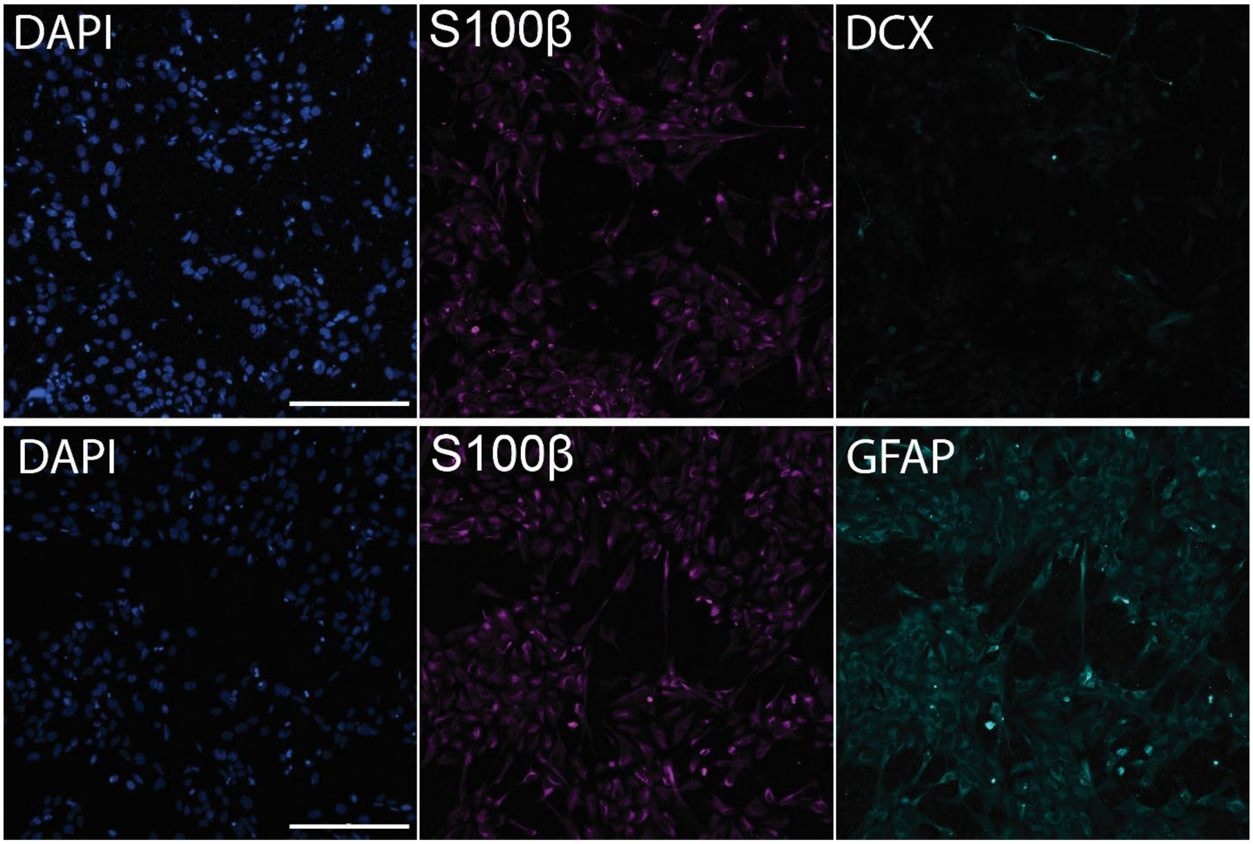

**Figure S3. Validation of astrocyte differentiation by immunocytochemistry.** Channels shown separately for image in **Fig. 3**. Scale bar = 200 μm

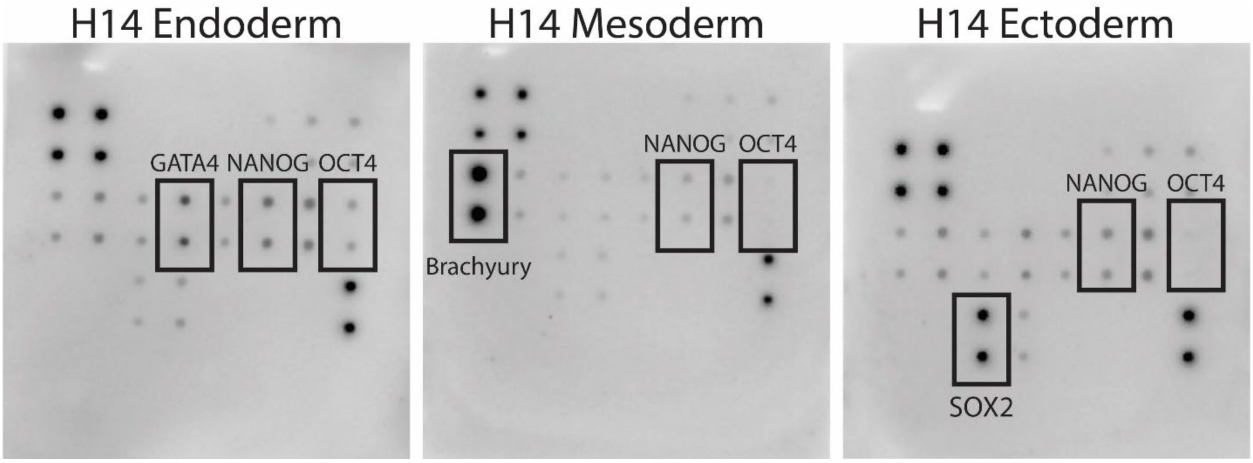


**Figure S4. Additional validation of stem cell pluripotency by trilineage differentiation.**

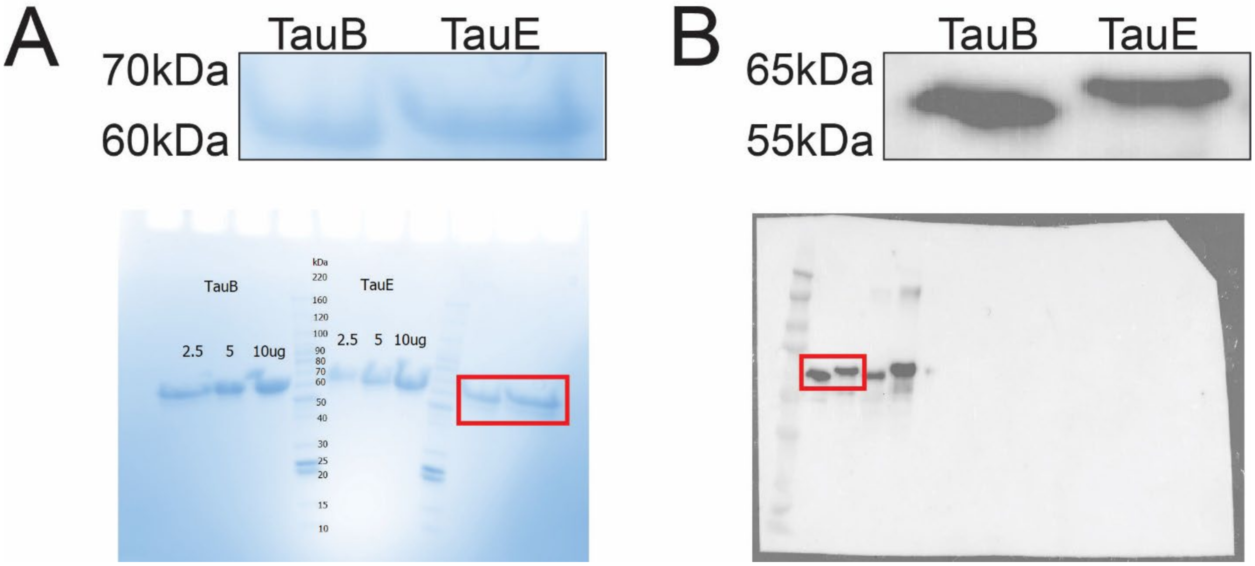


**Figure S5. Validation of recombinant tau.** (A) Recombinant TauE (1N4R) and TauB (1N3R) was run on an SDS-PAGE gel and stained with Coomassie Blue to assess for purity. Red box on whole gel represents magnified region shown above. (B) TauE and TauB were run on an SDS-PAGE gel, transferred to PVDF membrane and probed with anti-tau (HT7). Red box indicates relevant area magnified above

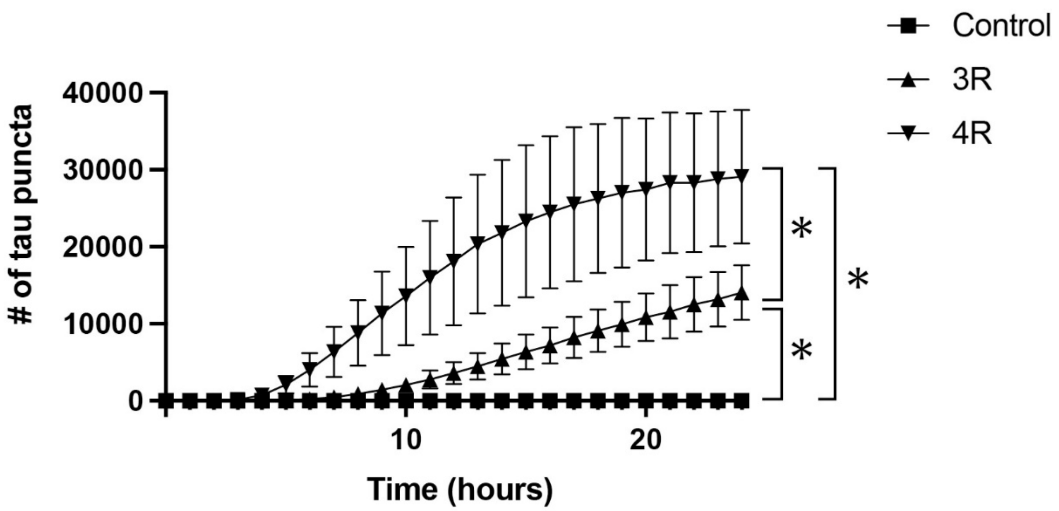


**Figure S6. Replication of Figure 3**. N=3 technical replicates per condition; *p<10-4

**Table S1. List of all genes for Figure 4.**
